# Cpipe: a shared variant detection pipeline designed for diagnostic settings

**DOI:** 10.1101/020388

**Authors:** Simon P Sadedin, Harriet Dashnow, Paul A James, Melanie Bahlo, Denis C Bauer, Andrew Lonie, Sebastian Lunke, Ivan Macciocca, Jason P Ross, Kirby R Siemering, Zornitza Stark, Susan M White, Melbourne Genomics Health Alliance, Graham Taylor, Clara Gaff, Alicia Oshlack, Natalie P Thorne

## Abstract

The benefits of implementing high throughput sequencing in the clinic are quickly becoming apparent. However, few freely available bioinformatics pipelines have been built from the ground up with clinical genomics in mind. Here we present Cpipe, a pipeline designed specifically for clinical genetic disease diagnostics. Cpipe was developed by the Melbourne Genomics Health Alliance, an Australian initiative to promote common approaches to genomics across healthcare institutions. As such, Cpipe has been designed to provide fast, effective and reproducible analysis, while also being highly flexible and customisable to meet the individual needs of diverse clinical settings. Cpipe is being shared with the clinical sequencing community as an open source project and is available at http://cpipeline.org.

## Background

Diagnostic laboratories are rapidly adopting high throughput genomic sequencing for clinical genetic tests. This transition is enabling a dramatic expansion in our ability to diagnose and screen heterogeneous monogenic disorders [1]. One critical aspect of a clinical genomics test is the bioinformatics pipeline used to analyse the sequencing data and output variants for clinical consideration. Thus far most clinical sequencing analysis pipelines have been driven by individual laboratories, who have either developed their own bioinformatics capability for processing data, relied on commercial products, or have partnered with research institutions to acquire the expertise needed. This approach has enabled rapid adoption, but has resulted in a wide diversity of implementation approaches and great variability in the methods used for evaluation, interpretation and reporting of variants. When pipelines have been primarily developed for research use they often lack the robustness, provenance and quality control features, maintainability and high degree of automation required in the clinical diagnostic setting. Additionally, many such analysis pipelines are designed without prioritising the ability to generalise to different diseases, technologies or computational contexts. Commercial pipelines can address some of these problems. However they are inevitably constrained in the level of customisation and transparency they can offer due to their commercial nature. Additionally commercial pipelines can be expensive for laboratories to acquire, evaluate and deploy. Altogether these issues impede the standardisation of bioinformatics pipelines for routine diagnostics across multiple clinics and healthcare systems. An analysis pipeline that is specifically designed for the clinical setting and that can be informed and iteratively improved by the clinical diagnostic community has the potential to offer the most effective diagnostic value.

Recognising these issues, the Melbourne Genomics Health Alliance was formed as a collaboration between seven institutions, including hospitals, diagnostic laboratories, universities and research institutes, with the aim of developing a common approach to the analysis and management of genomic data within Australia’s publicly funded healthcare system. A key outcome of the Alliance has been the development of a consensus bioinformatics pipeline, which we have called Cpipe. Cpipe is founded upon best practice analysis components that are emerging in the global clinical sequencing community and are already being employed by many of the members of the Alliance. However, the goal of Cpipe is not to improve upon these core bioinformatics analysis methods, nor is it ultimately to focus on any particular tool set. Rather, the aim of Cpipe is to create a common framework for applying the tools, that can be readily adapted for a diverse range of diagnostic settings and clinical indications.

We identified three key requirements for a clinical bioinformatics pipeline that differ from a pipeline intended for research use. First, a clinical pipeline must be designed with a greater emphasis on robust and reproducible analysis. There must be clear records of what analysis was performed and what files were used to generate results. Second, a number of specialised bioinformatics steps are required in clinical settings. For example, one key difference in a clinical setting is the need for variants to be assessed for their relevance to a given patient. Therefore it becomes vital to filter and prioritise variants to speed up this process and thus reduce the time clinicians spend assessing variants. Finally, the pipeline must be highly transparent and modular, so that the individual steps as well as the overall flow of the pipeline are easy to understand and modify. These qualities are critical in the clinical environment to allow laboratories to maintain and adapt pipelines to their needs without compromising on quality.

There have been a number of previous efforts to create publicly available analysis pipelines for high throughput sequencing data. Examples include Omics-Pipe [2], bcbio-nextgen [3], TREVA [4] and NGSane [5]. These pipelines offer a comprehensive, automated process that can analyse raw sequencing reads and produce annotated variant calls. However, the main audience for these pipelines is the research community. Consequently, there are many features required by clinical pipelines that these examples do not fully address. Other groups have focused on improving specific features of clinical pipelines. The Churchill pipeline [6] uses specialised techniques to achieve high performance, while maintaining reproducibility and accuracy. However it is not freely available to clinical centres and it does not try to improve broader clinical aspects such as detailed quality assurance reports, robustness, reports, and specialised variant filtering. The Mercury pipeline [7] offers a comprehensive system that addresses many clinical needs: it uses an automated workflow system (Valence, [8]) to ensure robustness, abstract computational resources, and simplify customisation of the pipeline. Mercury also includes detailed coverage reports provided by ExCID [9], and supports compliance with US privacy laws (HIPAA) when run on DNANexus, a cloud computing platform specialised for biomedical users. Mercury offers a comprehensive solution for clinical users, however it does not achieve our desired level of transparency, modularity and simplicity in the pipeline specification and design. Further, Mercury does not perform specialised variant filtering and prioritisation that is specifically tuned to the needs of clinical users.

Cpipe focuses on implementing or improving the three key aspects of clinical analysis pipelines that we have identified. The first aspect includes features that support the robustness and quality of the pipeline operation and these are provided automatically in Cpipe by the underlying pipeline framework, Bpipe [10]. The second aspect is the addition of specialised bioinformatics steps that are required for clinical settings. These include detailed quality reports, additional filtering and prioritisation of variants, and carefully designed output formats that accelerate clinical interpretation. Finally, Cpipe aims to be highly transparent and modular, so that it is easy to understand and modify the underlying tools used. This is critical to ensuring that Cpipe can be deployed in diverse clinical settings and can be updated and shared between different organisations, while still maintaining a common underlying framework.

Cpipe has been developed in close consultation with many different stakeholders from the clinical and research sequencing community in Melbourne, Australia. It is being actively used by three separate institutions for clinical sequencing, and is undergoing accreditation for diagnostic use. By adopting Cpipe, a solution that has already been tested in a diagnostic context, a laboratory can save significant effort in developing a pipeline. Perhaps even more importantly, by adopting Cpipe they can become part of a community of users and developers, and can benefit from the ongoing maintenance and active development that will occur over time. The open source license of Cpipe (GPLv3) will allow users of Cpipe to become contributors to the project, further ensuring its ongoing maintenance and development.

## Implementation

### Cpipe is built using Bpipe

Cpipe is implemented using a pipeline construction framework called Bpipe [10]. Bpipe automatically provides many features supporting our goals in creating Cpipe. Bpipe and its features are central to our implementation. Therefore we named the pipeline Cpipe, emphasising the close relationship between the two, and with the “C” indicating the clinical nature of the pipeline.

One of the most notable features of Bpipe is its pipeline construction language, which allows commands to be specified in a form that is nearly identical to executing them manually. This greatly increases the accessibility of Bpipe pipelines, as users do not need to learn a specific programming language or use specialised syntax to understand existing pipelines or to make simple modifications. Another powerful feature of Bpipe is that it automatically adds robustness features to every command executed with minimal intervention from the user. These features include automatic tracking of command history, logging of input and output files, clean-up of partially created files from failed commands, dependency tracking, automatic removal of intermediate results, generation of graphical reports, tracking of performance statistics, and notifications by email and instant messaging in response to failures. The audit trail created by this process can be used to reproduce or verify any part of any previous analysis.

Another key feature that Bpipe offers is abstraction from the computational environment. That is, Bpipe enables the same pipeline to easily work on a computational cluster, a local server or even a stand-alone desktop computer. This feature is important for building a pipeline that can be deployed in many different environments. To facilitate maximum utilisation of resources, Bpipe supports parallelisation, so that independent steps can be run simultaneously with minimal effort from the user. These features enable Cpipe to utilise cluster infrastructure where available, but importantly, Cpipe can automatically adapt to environments where significant parallelisation is not an option. Cpipe parallelises firstly by aligning reads from each lane and sample in parallel. After the initial alignment, processing is parallelised only by sample, and by parallelising selected independent operations at the sample level.

Generation of reports and evidence about the operation of the pipeline is a key requirement in clinical settings. Bpipe offers built in template-driven report generation features. These operational reports can be easily and automatically attached to emails that are sent in response to events that occur as part of the analysis. This makes it possible for operators to be alerted by email when pipeline errors or QC issues occur. A final important aspect of Bpipe is the high-level job management capabilities. Bpipe gives the operator the ability to start a pipeline with a single command, and to easily stop or view the status of running pipelines.

### Cpipe Architecture

#### Analysis Profiles

At the root of Cpipe’s architecture is the assumption that, in a clinical diagnostic setting, sequencing runs will be performed on many different patients, each of whom may have a different disease. These different diseases may require not only differing genes to be prioritised, but also different settings or tools to be applied in the analysis pipeline. As the field matures, it is even likely that patients with the same disease will be prescribed personalised diagnostic tests based on their individual phenotypes. However, this variability presents challenges, because most pipelines use a single set of targeted genes and tool settings for all samples in the analysis. To address this problem, Cpipe defines the concept of an “analysis profile”. The analysis profile is predefined to optimise settings for a particular subgroup of patients, such as those with a common clinical diagnosis. A specific analysis profile is assigned to each sample as an input to the pipeline. The parameters defined in the analysis profile can include: the list of genes to be included or excluded in the analysis; minimum quality and coverage thresholds for variants that are reported; the width of the window beyond exonic boundaries that should be used to identify potential splice site variants; and any other customisable settings that could be applicable to different patients. Cpipe supports definition of new customisable settings in a simple manner via a text file that can be supplied as part of the analysis profile definition for each sample. By using fixed, predefined, analysis profiles, laboratories can validate and accredit each profile independently as the need arises. This strikes a balance between customisation for each sample and the needs of accreditation agencies to have tests validated in advance. In the context of the Melbourne Genomics Health Alliance, the same exome capture platform was used for every patient but distinct gene sets were reported depending on the phenotype of the patient.

#### Directory Structure

Cpipe defines a standard directory structure that is used for all analyses. This predefined structure has two important benefits. Firstly, it enhances maintainability and usability of the pipeline. Secondly, it ensures that operational parts of the pipeline are well separated from parts of the pipeline that should not be modified. For each analysis, all the inputs, outputs and design files are isolated in a single “batch” folder so that each batch is completely isolated from other batches (Figure 1). When an analysis runs for the first time, all of the files that are defined in the analysis profile are copied to a dedicated “design” folder so that if the analysis is re-executed in the future, the same results will be produced. These factors help to ensure the reproducibility of results.

**Figure 1.**
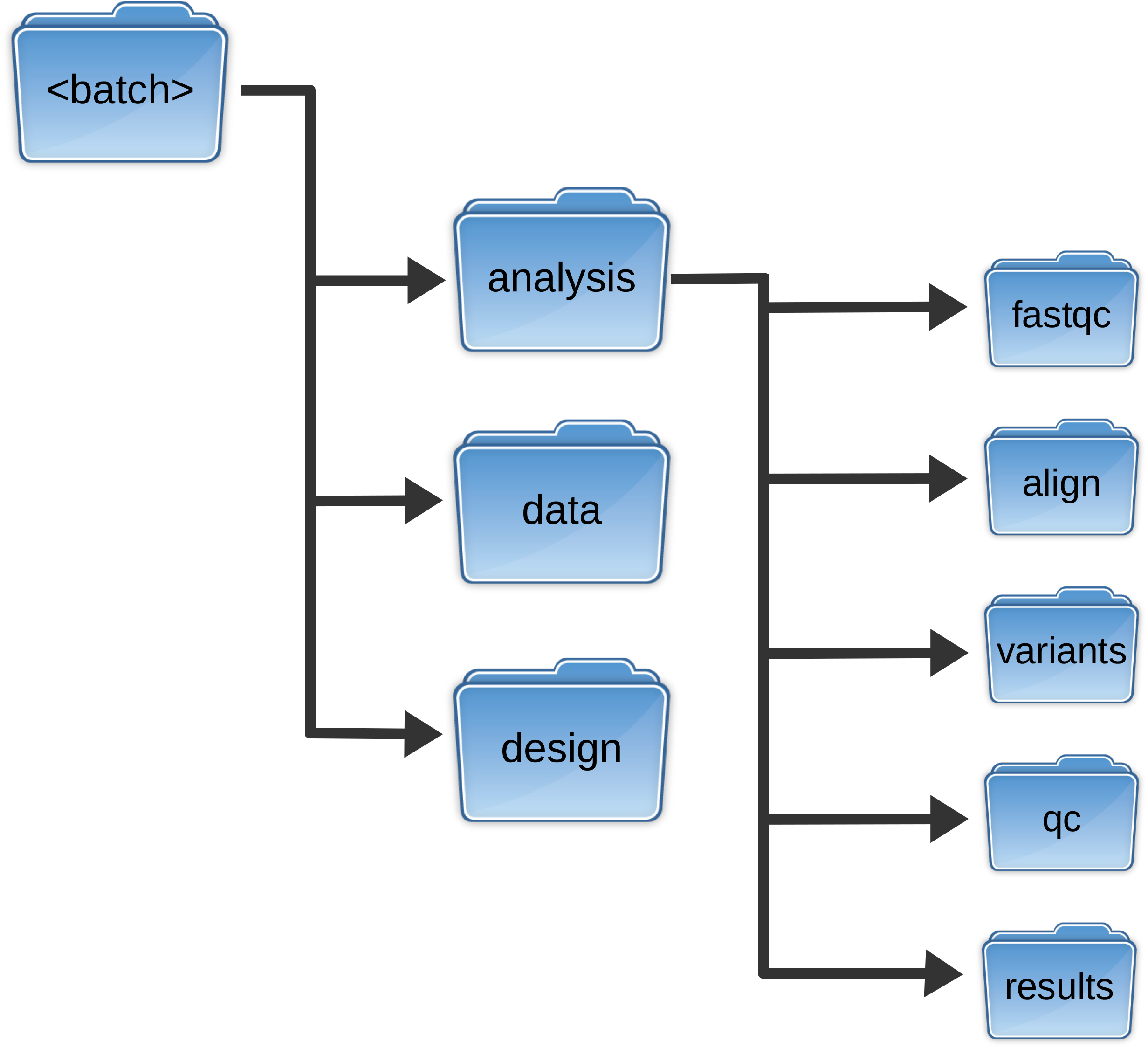
Batch directory structure used by Cpipe. Each analysis is conducted using a standardised directory structure that separates raw data, design files and generated results from each other. All computed results of the analysis are confined to the “analysis” directory, while source data is kept quarantined in the “data” directory. The analysis directory keeps separate directories for each stage of the analysis starting with initial quality control (fastqc), alignment (align), variant calling (variants) and final quality control (qc). The final analysis results are placed in the “results” directory.

#### Bioinformatics Analysis Process

The core bioinformatic analysis implemented by Cpipe (Figure 2) is based on the approach developed and recommended by the Broad Institute [11], and generally accepted by the community as best practice. This workflow includes: alignment using BWA mem [12], duplicate removal using Picard MarkDuplicates [13], Indel realignment using the GATK IndelRealigner, base quality score recalibration using the GATK BaseRecalibrator, and variant calling using the GATK HaplotypeCaller. The Broad Institute guidelines were developed for use in a research setting, and thus require some modifications for use in a clinical setting. Modifications in Cpipe include: (1) using Annovar [14] for annotation of variants as this tool provided a more comprehensive set of annotations desired by the clinical users in the Melbourne Genomics Health Alliance; (2) calling variants in each sample separately instead of using joint calling, as this ensures that results for a sample can be reproduced without requiring data belonging to other samples; (3) no variant quality score recalibration is performed because variant quality scores themselves are not used in downstream filtering by Cpipe, and because unless a large independent reference sample set is created, the procedure causes inter-sample dependencies.

**Figure 2.**
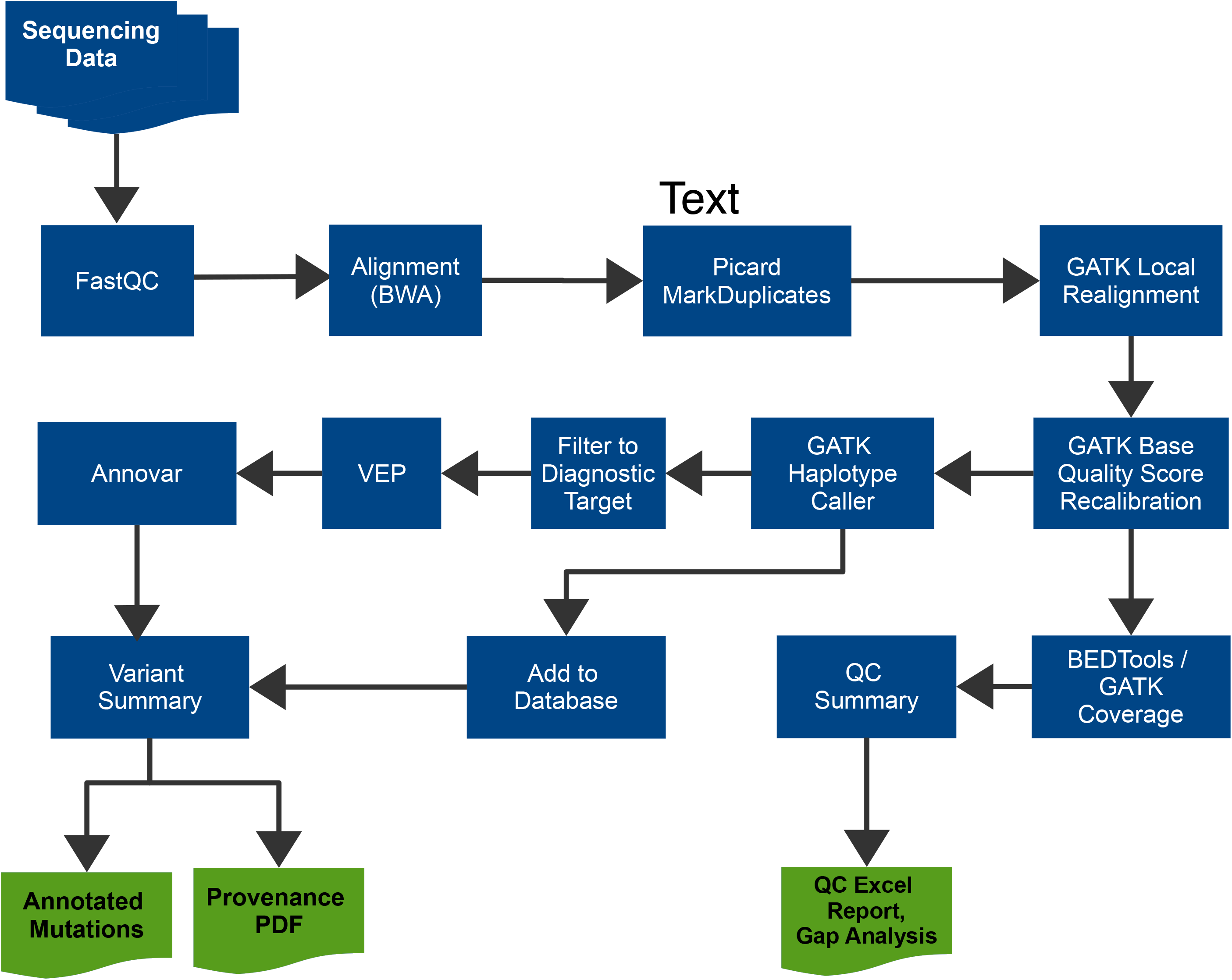
Simplified Cpipe analysis steps. Cpipe consists of a number of steps. The core of these are based on the best practice guidelines published by the Broad Institute, consisting of alignment using BWA mem, duplicate removal using Picard MarkDuplicates, local realignment and base quality score recalibration using GATK, and variant calling using GATK HaplotypeCaller. To support clinical requirements, many steps are added including quality control steps (BEDTools coverage and QC summary), additional annotation (Annovar and the Variant Effect Predictor, VEP) and enhanced reports (Annotated variants, Provenance PDF, QC Excel report & Gap Analysis).

The analysis process described in this section makes use of two components (GATK and Annovar) that may require a license for clinical use. To allow use of the pipeline without licensing these components, Cpipe supports alternative options. To substitute for GATK, Cpipe allows use of an older version of GATK that is free to use commercially. The Variant Effect Predictor and SnpEFF [15] are supported as alternative options to Annovar that are free for commercial and clinical use.

The default pipeline that Cpipe implements is designed as a sound baseline that caters to a broad set of clinical needs. However it is fully intended that laboratories will tune these components and potentially replace them with different tools that may be better suited to a particular application. The current default Cpipe workflow is intended for analysis of single, unrelated samples. Analysis of related samples requires joint calling within each family to provide fully informative results. This feature is currently being implemented and will be released in a future version of Cpipe.

#### Internal Variant Database

A common diagnostic strategy for rare diseases is to filter out variants that are observed at a frequency in the population that is inconsistent with the prevalence of the disease. High throughput sequencing typically identifies many thousands of variants that are observed in multiple samples. These variants are often not present in public population databases either due to them being population-specific or technical artefacts. Cpipe therefore maintains an internal database of all variants observed in all samples that are processed by that specific instance of Cpipe. The frequency of observations in this internal database may be used as a criterion for excluding variants, alongside allele frequencies annotated from public databases. The internal database is implemented using SQLite [16]. SQLite is a fully embedded database technology that stores all data in a single, stand-alone file. This simplifies the configuration and installation of the pipeline by removing the need for an external database server.

The internal variant database accumulates variants over time as more analyses are run. Therefore, a sample that is re-analysed by Cpipe at a later date may be assigned different values for the frequency at which variants are observed in the internal database. Apart from this single measure, however, Cpipe is designed so that entering identical input data always produces identical analysis results. To ensure complete reproducibility, the SQLite database file may be archived to capture a snapshot of the database prior to each analysis.

#### Quality Control Reports

In the diagnostic setting, it is critical to assess which regions of a gene were adequately interrogated by the test, so that clinicians can determine if additional sequencing is required to detect a causative variant in that gene. It is therefore necessary that detailed information about sequencing coverage is provided in QC reports. Cpipe supports this requirement by producing three separate reports: a gene level report, an exon level report and a detailed base-pair level gap report. These allow a curator or clinician to rapidly understand, at a high level, the quality of the sequencing coverage, and then to investigate in more detail if a particular gene or exon is of concern.

The scale of clinical operations means that only a small number of staff may be responsible for running many simultaneous analyses. It is therefore important that as many essential quality checks as possible are automated. Cpipe uses the Bpipe ‘check’ feature to support automated checks in the pipeline. Failure of these checks results in an automated email notification to the pipeline operator with an attached document describing the failure. These include: (i) failure of a sample if specific FASTQC measures fail, (ii) failure of a sample if the overall median coverage falls below a configurable threshold, (iii) failure if the median fragment size of the sequenced reads falls outside a user configurable range, (iv) failure of a sample if the rate of PCR duplicates is greater than a user configurable threshold, (v) failure of a sample if a bioinformatic check of the sex of the sample is inconsistent with the sex declared for the sample in the inputs to the pipeline.

#### Prioritisation, Categorisation and Filtering of Variants

One of the most significant challenges in bringing high throughput sequencing into routine clinical care is that of scaling the difficult and highly manual job of curation, classification/interpretation and reporting of variants. This task frequently presents a ‘bottleneck’ in diagnostic workflows, limited by the number of trained staff with the required expertise to evaluate the variants and report the results. To address this, Cpipe implements a filtering and prioritisation system designed to automatically highlight the results most likely to be relevant for the majority of cases. This system was designed in close collaboration with clinicians in the Melbourne Genomics Health Alliance and aims to reflect the usual approach taken by a curator when first faced with a variant list from a given patient. The approach consists of two strategies that dramatically reduce the number of variants to be clinically considered in the first instance (Figure 3).

**Figure 3.**
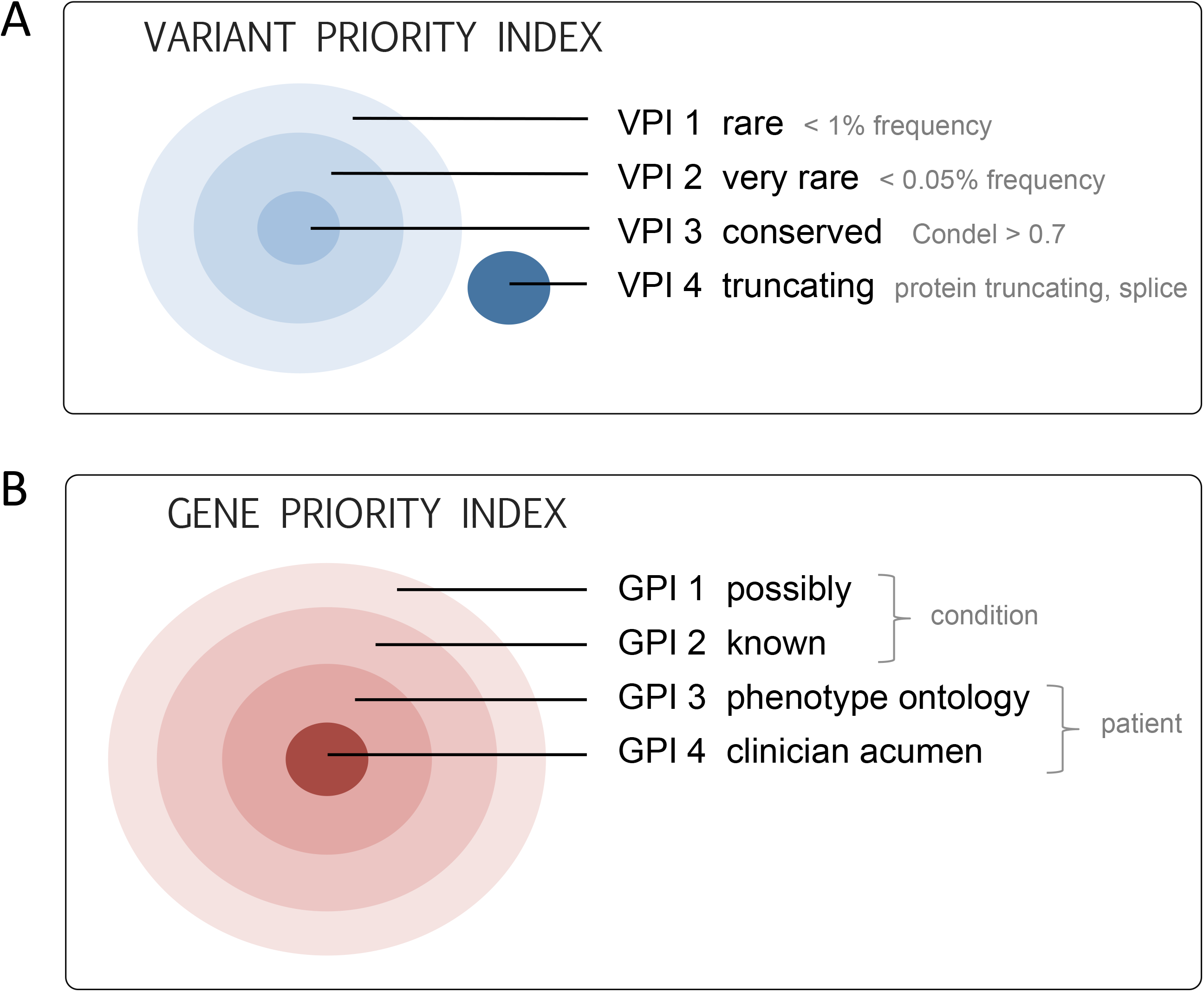
Variant and Gene Priority Indexes. Curation of variants is aided by a prioritisation system that ranks variants according to (A) characteristics of the variant including frequency in population databases, conservation scores and the predicted impact on protein product, and (B) the strength of association of the gene to the phenotype under consideration.

The first is a specifically defined system, the Variant Priority Index, that combines a range of factors to place variants into four distinct tiers (Figure 3A). The tiers are ordered according to measures of rarity, conservation and truncating effect on the transcript protein. Tiers one, two and three are subsets of each other. Tier one (VPI 1) corresponds to “rare” in-frame indels or missense variants with frequency less than 0.01 in EVS [17], 1000G [18] and ExAC [19]. Variants are elevated to tier two (VPI 2) “very rare or novel” if their frequency in these population databases is less than 0.0005. Likewise, tier two variants are promoted to tier three (VPI 3) if they are also “highly conserved” (Condel>0.07)[20] as well as “very rare or novel”. VPI 4 is reserved for the highest priority variants including frameshift, truncating, and splice site variants. The tiers provide an intuitive first pass prioritisation of variants, making it easier for curators to quickly see potentially important variants and therefore helping to manage their workload. Variants that do not meet the criteria for at least VPI 1, are hidden in the result set.

The second strategy is a prioritisation of genes into categories based on a-priori likelihoods for being causal to the specific patient (Figure 3B). The Gene Prioritisation Index starts with all genes in the analysis profile target region (GPI 1), then narrows to genes that are commonly known to be causal for the disease or patient group (GPI 2), and finally narrows again to a set of custom genes that may be prioritised by the patient’s clinician based on individual considerations, such as phenotype, using either insilico programs (GPI 3) or their own clinical acumen (GPI 4).

#### Output Results

The final result of the bioinformatics pipeline is a spreadsheet containing filtered and annotated variants. The format of this spreadsheet is designed to aid rapid interpretation by curators. Variants are sorted by the previously described Variant Priority Index and Gene Priority Index such that the most promising variants are sorted to the top of the spreadsheet.

As an adjunct, a set of files in CSV format is produced that contain identical information to the spreadsheet, but which are formatted in such a way as to facilitate input into an LOVD3 [21] compatible database. Exploiting this capability, the Melbourne Genomics Health Alliance has developed an enhanced version of LOVD3 (MG-LOVD) that includes functionality to greatly facilitate the curation, classification/interpretation and reporting process (paper in preparation).

#### Regression Tests

All aspects of the technology surrounding clinical genomics are quickly evolving. It is thus essential that software pipelines are readily adaptable to new changes. However such changes must be validated to ensure they do not affect the clinical results of the pipeline in an unexpected way. To assist with this, Cpipe includes a set of automated software regression tests, which operate as a “self-test module”. The first of these tests analyses sequence data from chromosome 22 of the Coriell sample NA12878 [22], and then compares results to a set of predefined high confidence calls published by Illumina as part of the Platinum Genomes Project [23]. The test fails if insufficient sensitivity is observed. A second test simulates variants in data from the same sample using a simulation tool, Bamsurgeon [24], to test detection and correct annotation of a range of variants that would typically be treated as clinically significant. Finally, the self-test module performs a number of additional software regression tests to confirm that the automated quality checks in the pipeline are functioning correctly. These tests do not substitute for the full and rigorous validation required by accreditation agencies. However, they nonetheless play a vital role in supporting ongoing development by providing immediate feedback about the impact of any change on the pipeline.

### Results and Discussion

We have implemented Cpipe, an exome analysis pipeline designed specifically for the needs of clinical users. Cpipe has been developed through an extensive process of consultation between many different stakeholders involved in the Melbourne Genomics Health Alliance including bioinformaticians, IT specialists, sequencing laboratories, diagnostic users and genetic and specialist clinicians. Cpipe takes raw sequence data and patient specific analysis profiles and performs variant calling and prioritisation. In addition it provides multiple reports including QC reports and provenance files. Results of Cpipe can also be imported into public variant databases (Figure 4).

**Figure 4.**
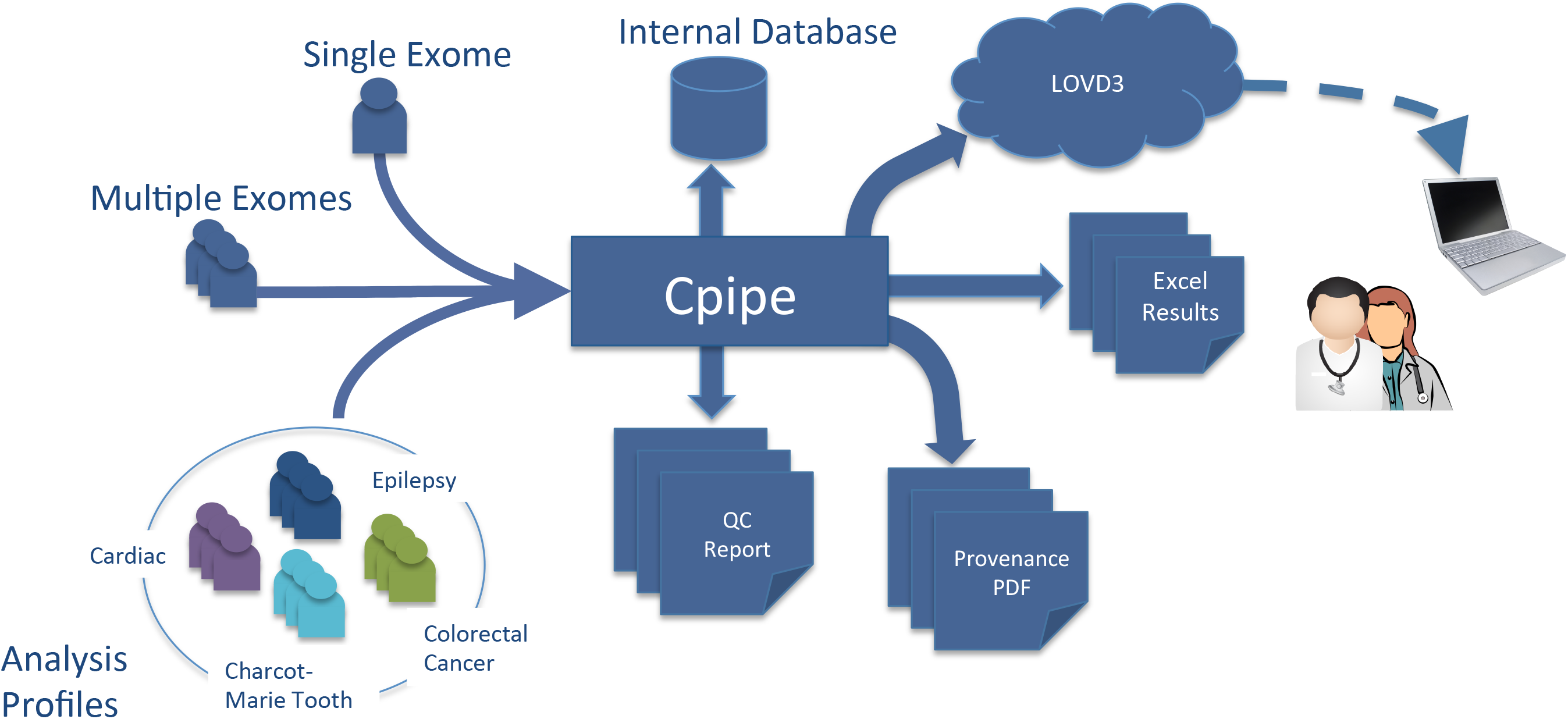
Overview of Cpipe workflow. Cpipe accepts a flexible arrangement of exome or targeted capture samples. Each sample is assigned an Analysis Profile that determines the particular settings and gene list to analyse for that sample. Provenance and QC reports are produced as Excel and PDF files, while variant calls are delivered as both an Excel spreadsheet and a CSV file that is importable to LOVD3. In addition to allele frequencies from population databases, allele frequencies are also annotated from an internal embedded database that automatically tracks local population variants and sequencing artefacts.

#### Evaluation in Production Setting

After development, Cpipe was deployed in an operational diagnostic setting and to date has been used to analyse 168 exomes as part of the Melbourne Genomics Health Alliance demonstration project. This project was designed to prototype the deployment of exome sequencing as a clinical diagnostic test within a health system in the states of Victoria and Tasmania in Australia. The samples were chosen from five diverse disease cohorts to evaluate different diagnostic applications. Results from Cpipe were imported into an instance of MG-LOVD database that was presented to curators and clinicians to facilitate the identification of causal variants for each patient. While the specific diagnostic outcomes for the Melbourne Genomics Health Alliance demonstration project will be reported elsewhere, we found that using the outputs generated by Cpipe, the diagnostic rate for a broad range of Mendelian adult and childhood conditions compares favourably to well established clinical genomics projects that claim diagnostic rates of 25%-35% [25, 26].

Samples were sequenced using Illumina HiSeq 2500 instruments after capture by the Nextera V1.2 exome capture kit. Sequencing was performed at two laboratories, the Australian Genome Research Facility and the Centre for Translational Pathology, The University of Melbourne. Samples were sequenced and processed in batches of 12, yielding approximately 50 million reads per sample. Median coverage depth for each sample varied between 75 and 254 (median=129, n=168). To process the samples, Cpipe was deployed on a 32 core system with 1TB of RAM and a high performance GPFS storage system. Typically, Cpipe processed a batch of 12 samples in 24 hours. On average each sample required a peak of 21 GB of space, however Cpipe automatically reclaims space used by intermediate files so that the mean space consumed per sample was 15 GB. While the processing time for an example batch of 12 samples was 24 hours, 28 minutes, the total computation time accumulated by all processing stages for 12 samples was approximately 187 hours. Bpipe’s automatic parallelisation features thus allowed significant reduction in the processing time.

#### Variant Prioritisation and Filtering

The combination of the Variant Prioritisation Index, Gene Prioritisation Index and filtering significantly reduces the number of variants prioritised for curation. For example, a small gene panel of 55 candidate genes yielded only 2 variants per sample to be curated on average (ranging from 0 to 6, n=31). For larger panels containing up to 3,000 genes, there were on average 115 variants left after filtering that required curation (ranging from 76 to 183, n=37). This was reduced to an average of 1.45 variants per patient (ranging from 0 to 6, n=35) when the treating clinician defined a group of genes as Gene Priority Index 4. The average number of genes in Gene Priority Index 4 was 21 (ranging from 1 to 100, n=35).

In the operational setting where the Melbourne Genomics Health Alliance has processed 168 samples, we observe that 89% of all non-synonymous coding variants are removed by filtering on allele frequency in the 1000 genomes project [18] and the Exome Sequencing Project [17]. As described, Cpipe also uses an internal variant database to filter out variants that are observed in multiple samples and that belong to different disease cohorts. A further 39% of the remaining variants were able to be removed by filtering using the internal variant database. This demonstrates that even after filtering using public databases, maintaining a local variant database is still important for removing common private population variants and artefacts introduced by sequencing or bioinformatic steps.

#### Variant Calling Performance

To check the variant calling performance achieved by Cpipe using the default GATK based tool set described earlier, reads from the 1000 Genomes sample NA12878 were analysed. This sample was sequenced to a median coverage depth of 91X as part of the Melbourne Genomics Health Alliance demonstration project. The resulting variant calls were compared to a set of high confidence calls obtained from the Illumina Platinum Genomes Project (Version 7.0) [23]. For regions in the exome target Cpipe achieved 90.2% sensitivity to SNVs in the high confidence set with a false discovery rate of 9%. After filtering the high confidence calls to include only regions where our sample had greater than 15x coverage, sensitivity increased to 95.7%. These rates are indicative of the default variant calling performance achieved by Cpipe. However we emphasise that Cpipe is a framework designed specifically to allow users to customise the individual tools to suit their needs. Thus different variant calling options, or an entirely different variant calling tool can be easily substituted to modify performance to the needs of a particular application.

#### QC Reports

We analysed the healthy control sample NA12878 for a gene panel previously published for diagnosis of cardiomyopathy patients [27] to generate examples of the QC reports generated by Cpipe. The gene report [see Addtional file 1] provides a high level view that allows a curator to quickly assess whether coverage is adequate over the genes of interest with a color-coded system. 2 out of 20 genes from the panel were identified as having potentially unsatisfactory coverage. The exon-level report details which exons within these genes of interest have insufficient coverage. In this case, 12 exons were reported as being only partially covered, representing 32% of the total exons in poor quality genes [see Additional file 2]. The gap report allows exact identification of all regions having coverage below a fixed, user-configurable threshold [see Additional file 2]. Thus a curator can discover at sub-exon level which regions have poor coverage and potentially suggest follow-up sequencing to address these specific genomic positions. Our test sample contained 55 distinct regions having poor coverage. These regions accounted for 1.3kb of sequence in total (3.8% of the gene panel target regions).

The built in QC reporting features provided by Cpipe allow clinical users to quickly and easily ascertain if sequencing has achieved sufficient quality to diagnose a patient. A feature of the Cpipe framework is that it is very straightforward to customise these reports and to add new reports.

## Conclusions

We have presented Cpipe, a new exome and targeted sequencing analysis pipeline that is designed specifically to support clinical needs. As clinical implementation of sequencing data becomes widespread there is a need for a freely available analysis platform that can be shared between clinical laboratories. Cpipe is currently in routine use at three separate institutions in Melbourne and is undergoing accreditation for diagnostic use. These organisations are actively maintaining the common pipeline. Cpipe is made available by the Melbourne Genomics Health Alliance under the open source GPLv3 license, allowing full and free use of the pipeline for both commercial and non-commercial purposes. By adopting Cpipe as their clinical sequencing pipeline framework, other members of the clinical sequencing community can benefit, not just from a pipeline that already contains many needed features, but also from the ongoing development that will occur over time.

## Availability and requirements

**Project Name**: Cpipe

**Project Home Page:** http://cpipeline.org

**Operating system(s):** Linux / Unix

**Programming language:** Mixed: Java, Groovy, Python, Bash

**Other requirements:** Reference data, Java 1.7+, Perl 5.10+, Python 2.7+

**License:** GPLv3

**Any restrictions to use by non-academics:** Two programs (GATK and Annovar) that are required for the full features of the software may require a license for commercial use. Cpipe can work with a reduced feature set without these tools.

## List of abbreviations

HIPAA: Health Insurance Portability and Accountability Act
1000G: 1000 Genomes Project [http://1000genomes.org]
ExAC: Exome Aggregation Consortium [http://exac.broadinstitute.org]
LOVD: Leiden Open Variation Database

## Authors’ contributions

SPS wrote the Cpipe software. HD made modifications, added support files and implemented processing of the data, PJ,NPT,IM,CG proposed the variant priority scheme, NPT,IM,ZS,SW,PJ,CG proposed the gene priority scheme,

SPS,HD,MB,DB,AL,IM,JR,KRS,GT,AO,NPT provided input into the structure and design of the pipeline, SPS wrote the majority of the paper, with significant contribution from HD,AO,NPT. CG, as the program leader, established the overall concept of Melbourne Genomics Health Alliance. AO supervised SPS and co-chaired the Genomics and Bioinformatics Advisory Group that advised on the pipeline development. NPT managed the development of this project.

## Acknowledgements

We thank Charlotte Anderson and John-Paul Plazzer for their contributions to discussions involving the interoperability of Cpipe with the Melbourne Genomics Health Alliance enhanced version of LOVD (MG-LOVD). We thank Melissa Martyn for her assistance in providing the evaluation data from the Melbourne Genomics Health Alliance demonstration project. This work has been supported by the Victorian Government’s Operational Infrastructure Support Program and Australian Government NHMRC IRIISS. MB was supported by an ARC Future Fellowship (FT100100764) and an NHMRC Program Grant (APP1102971). AGRF is supported by Commonwealth government infrastructure funding schemes administered through Bioplatforms Australia including the National Collaborative Research Infrastructure Scheme 2 (NCRIS 2).

## Additional Files

**Additional File 1**

An example of the sample summary PDF produced by Cpipe for high level quality control purposes. This example is produced from sequencing reads for 1000 Genomes sample NA12878 over a panel of genes related to cardiomyopathy.

**Additional File 2**

An example of the detailed quality control spreadsheet containing coverage gaps and exon level coverage details for each gene in the diagnostic target region of the analysis profile. This example is produced from sequencing reads for 1000 Genomes sample NA12878 over a panel of genes related to cardiomyopathy.

**Additional File 3**

An example of the final report in Excel format produced by Cpipe. This example is produced from sequencing reads for 1000 Genomes sample NA12878 over a panel of genes related to epilepsy.

## References

1 Rehm HL: Disease-targeted sequencing: a cornerstone in the clinic. Nat Rev Genet 2013, 14:295–300.

2 Fisch KM, Meissner T, Gioia L, Ducom J-C, Carland TM, Loguercio S, Su a. I: Omics Pipe: a community-based framework for reproducible multi-omics data analysis. Bioinformatics 2015(January):1–5.

3 bcbio-nextgen - Validated, scalable, community developed variant calling and RNA-seq analysis https://github.com/chapmanb/bcbio-nextgen, Accessed 31 March 2015.

4 Li J, Doyle M a., Saeed I, Wong SQ, Mar V, Goode DL, Caramia F, Doig K, Ryland GL, Thompson ER, Hunter SM, Halgamuge SK, Ellul J, Dobrovic A, Campbell IG, Papenfuss AT, McArthur G a., Tothill RW: Bioinformatics pipelines for targeted resequencing and whole-exome sequencing of human and mouse genomes: A virtual appliance approach for instant deployment. PLoS One 2014, 9.

5 Buske F a., French HJ, Smith M a., Clark SJ, Bauer DC: NGSANE: A lightweight production informatics framework for high-throughput data analysis. Bioinformatics 2014, 30:1471–1472.

6 Kelly BJ, Fitch JR, Hu Y, Corsmeier DJ, Zhong H, Wetzel AN, Nordquist RD, Newsom DL, White P: Churchill: an ultra-fast, deterministic, highly scalable and balanced parallelization strategy for the discovery of human genetic variation in clinical and population-scale genomics. Genome Biol 2015, 16:6.

7 Reid JG, Carroll A, Veeraraghavan N, Dahdouli M, Sundquist A, English A, Bainbridge M, White S, Salerno W, Buhay C, Yu F, Muzny D, Daly R, Duyk G, Gibbs RA, Boerwinkle E: Launching genomics into the cloud: deployment of Mercury, a next generation sequence analysis pipeline. BMC Bioinformatics 2014, 15:1–11.

8 Valence - Fast, Dynamic, and Extendable/Mergeable Workflow Management System. http://sourceforge.net/projects/valence/. Accessed 31 March 2015.

9 The Exome Coverage and Identification Report. https://github.com/cbuhay/ExCID. Accessed 31 March 2015.

10 Sadedin SP, Pope B, Oshlack A: Bpipe: a tool for running and managing bioinformatics pipelines. Bioinformatics 2012, 28:1525–6.

11 GATK Best Practices. https://www.broadinstitute.org/gatk/guide/best-practices?bpm=DNAseq. Accessed 31 March 2015.

12 Li H: Aligning sequence reads, clone sequences and assembly contigs with BWA-MEM. 2013:3.

13 Picard: A set of Java command line tools for manipulating high-throughput sequencing data (HTS) data and formats https://broadinstitute.github.io/picard/ Accessed 21 May 2015.

14 Wang K, Li M, Hakonarson H: ANNOVAR: functional annotation of genetic variants from high-throughput sequencing data. Nucleic Acids Res 2010, 38:e164.

15 Cingolani P, Platts A, Wang LL, Coon M, Nguyen T, Wang L, Land SJ, Lu X, Ruden DM: A program for annotating and predicting the effects of single nucleotide polymorphisms, SnpEff: SNPs in the genome of Drosophila melanogaster strain w 1118; iso-2; iso-3. Fly (Austin) 2012, 6:80–92.

16 Exome Variant Server, NHLBI GO Exome Sequencing Project (ESP), Seattle, WA [http://evs.gs.washington.edu/EVS/]

17 Durbin RM, Altshuler DL, Durbin RM, Abecasis GR, Bentley DR, Chakravarti A, Clark AG, Collins FS, Vega D La, Francisco M, Donnelly P, Egholm M, Flicek P, Gabriel SB, Gibbs R a., Knoppers BM, Lander ES, Lehrach H, Mardis ER, McVean G a., Nickerson D a., Peltonen L, Schafer AJ, Sherry ST, Wang J, Wilson RK, Gibbs R a., Deiros D, Metzker M, Muzny D, et al.: A map of human genome variation from population-scale sequencing. 2010.

18 Exome Aggregation Consortium (ExAC), Cambridge, MA. http://exac.broadinstitute.org. Accessed 31 March 2015.

19 González-Pérez A, López-Bigas N: Improving the assessment of the outcome of nonsynonymous SNVs with a consensus deleteriousness score, Condel. Am J Hum Genet 2011, 88:440–9.

20 Fokkema IFAC., Taschner PEM, Schaafsma GCP, Celli J, Laros JFJ, den Dunnen JT: LOVD v.2.0: The next generation in gene variant databases. Hum Mutat 2011, 32:557–563.

21 Coriell Institute for Medical Research. http://www.corriell.org. Accessed 31 March 2015.

22 Illumina Platinum Genomes. http://www.illumina.com/platinumgenomes/. Accessed 31 March 2015.

23 Ewing, A: Bamsurgeon. https://github.com/adamewing/bamsurgeon. Accessed 31 March 2015.

24 Yang Y, Muzny DM, Reid JG, Bainbridge MN, Willis A, Ward P a, Braxton A, Beuten J, Xia F, Niu Z, Hardison M, Person R, Bekheirnia MR, Leduc MS, Kirby A, Pham P, Scull J, Wang M, Ding Y, Plon SE, Lupski JR, Beaudet AL, Gibbs R a, Eng CM: Clinical whole-exome sequencing for the diagnosis of mendelian disorders. N Engl J Med 2013, 369:1502–11.

25 Wright CF, Fitzgerald TW, Jones WD, Clayton S, McRae JF, van Kogelenberg M, King D a, Ambridge K, Barrett DM, Bayzetinova T, Bevan a P, Bragin E, Chatzimichali E a, Gribble S, Jones P, Krishnappa N, Mason LE, Miller R, Morley KI, Parthiban V, Prigmore E, Rajan D, Sifrim A, Swaminathan GJ, Tivey AR, Middleton A, Parker M, Carter NP, Barrett JC, Hurles ME, et al.: Genetic diagnosis of developmental disorders in the DDD study: a scalable analysis of genome-wide research data. Lancet 2015, 385:1305–1314.

26 Gowrisankar S, Lerner-Ellis JP, Cox S, White ET, Manion M, LeVan K, Liu J, Farwell LM, Iartchouk O, Rehm HL, Funke BH: Evaluation of second-generation sequencing of 19 dilated cardiomyopathy genes for clinical applications. J Mol Diagn 2010, 12:818–827.

